# Identification of CD44 as a key mediator of cell traction force generation in hyaluronic acid-rich extracellular matrices

**DOI:** 10.1101/2023.10.24.563860

**Authors:** Brian C. H. Cheung, Xingyu Chen, Hannah J. Davis, Cassidy S. Nordmann, Joshua Toth, Louis Hodgson, Jeffrey E. Segall, Vivek B. Shenoy, Mingming Wu

**Affiliations:** Department of Biological and Environmental Engineering, Cornell University, Ithaca, NY, USA; Center of Engineering MechanoBiology, University of Pennsylvania, Philadelphia, PA, USA; Department of Materials Science and Engineering, University of Pennsylvania, Philadelphia, PA, USA; Department of Biological Sciences, Cornell University, Ithaca, NY, USA; Department of Biomedical Engineering, Cornell University, Ithaca, NY, USA; Department of Molecular Pharmacology, Albert Einstein College of Medicine, Bronx, NY, USA; Department of Pathology, Albert Einstein College of Medicine, Bronx, NY, USA

**Keywords:** 3D cell traction force microscopy, hyaluronic acid, CD44, collagen, tumor invasion

## Abstract

Mechanical properties of the extracellular matrix (ECM) critically regulate a number of important cell functions including growth, differentiation and migration. Type I collagen and glycosaminoglycans (GAGs) are two primary components of ECMs that contribute to mammalian tissue mechanics, with the collagen fiber network sustaining tension, and GAGs withstanding compression. The architecture and stiffness of the collagen network are known to be important for cell-ECM mechanical interactions via integrin cell surface adhesion receptors. In contrast, studies of GAGs in modulating cell-ECM interactions are limited. Here, we present experimental studies on the roles of hyaluronic acid (HA, an unsulfated GAG) in single tumor cell traction force generation using a recently developed 3D cell traction force microscopy method. Our work reveals that CD44, a cell surface adhesion receptor to HA, is engaged in cell traction force generation in conjunction with β1-integrin. We find that HA significantly modifies the architecture and mechanics of the collagen fiber network, decreasing tumor cells’ propensity to remodel the collagen network, attenuating traction force generation, transmission distance, and tumor invasion. Our findings point to a novel role for CD44 in traction force generation, which can be a potential therapeutic target for diseases involving HA rich ECMs such as breast cancer and glioblastoma.

## Introduction

The reciprocal mechanical interactions between mammalian cells and extracellular matrices (ECM) critically regulate a number of important physiological (e.g. immune response, wound healing) and pathological (e.g. fibrosis and cancer) processes(1-5). An important mediator of cell-ECM interaction is the cell traction force. Mammalian cells can form mechanical links to the surrounding ECM fibers using surface adhesion receptors, and then pull on the fiber network via traction force, a coordinated polymerization and reorganization of actin filaments through the action of myosin II(6). The cell traction force aligns the ECM fibers, which in return, can influence cell functions(5, 7). Cell-ECM mechanical interactions also provide a route for cells to communicate with each other via the mechanical tension generated within the ECM(8-11). In human breast carcinoma, it has been reported that tumor malignancy can be driven by ECM stiffness(3), and aligned collagen fibers perpendicular to the tumor boundary were found to correlate with poor survival(4).

Structurally, the ECM is a dynamic meshwork containing collagen fibers and glycosaminoglycans (GAGs). Mechanically, collagen sustains tension within the tissue and GAGs resist compression. Although extensive work has been done on how collagen networks regulate cell functions(8, 12-17), much less is known about the roles of GAGs in regulating cell-ECM interaction and subsequently cell function. Recently, theoretical work from the Shenoy lab has demonstrated that GAGs critically modulate ECM mechanical properties, as well as cell traction force generation transmission distance. Specifically, the presence of GAGs decreased the cell traction force transmission distance(18) and inhibited collagen alignment(18, 19). However, the role of GAGs in cell traction force generation has not been investigated experimentally.

Cell surface adhesion receptors and cytoskeletal molecules are two critical components for traction force generation(20-22). β1-integrins mechanically link the actin filaments within the cell with the collagen fibers within the ECM, and contribute to the traction force transmission from the cell to the ECM(23-27). In HA-rich ECMs, CD44, the adhesion receptor to HA(28, 29), is highly expressed in many cell types. Clinically, CD44 is associated with low survival in high grade breast cancer(30-34). In glioblastoma, it was demonstrated that CD44, but not integrin, is essential for HA-rich matrix stiffness sensing and cell migration(35, 36). Inspired by these observations, we asked whether or how CD44 contributes to traction force generation by providing a link between the ECM and the cytoskeletal molecules in HA-rich matrices. In this paper, we report our work on traction force measurements of breast tumor cells embedded within collagen and collagen-HA cogels. Our work reveals that CD44, in addition to β1-integrin, is required for cell force generation, and cell-induced ECM architectural remodeling, cell traction force and tumor invasion are all attenuated by the presence of HA.

## Results and Discussion

### Hyaluronic acid modulates the bulk mechanical properties and architecture of collagen-HA cogels

While normal breast tissues typically contain 0.1mg/mL HA, a 2-60 fold increase of HA content has been detected in various malignant tumors, and can result in a range of HA concentrations from about 0.2-6mg/mL(37, 38). To understand how HA alters the mechanics of the tumor microenvironment, we engineered the following three ECMs: i) 1.5mg/mL type I collagen (Col), ii) Col-HA cogel with 1.5mg/mL collagen and 1.5mg/mL HMW HA (H1), and iii) Col-HA cogel with 1.5 mg/mL collagen and 3.0mg/mL HMW HA (H2).

The microarchitecture of the ECMs was characterized using confocal microscopy in reflectance mode as well as scanning electron microscopy followed by a porosimetric analysis (Fig. 1A-C). By introducing HA into the collagen matrix, the collagen fiber networks of the H1 and H2 cogels have smaller pores and thinner fibers, compared to the collagen only gel. Using a rheometer, we investigated how these architectural changes within the collagen network translate into the bulk mechanical properties of the matrices (Fig. S1). Dynamic stress strain measurements at 0.1Hz using a parallel-plate rheometer were carried out at a range of strains. An example is shown in Fig. 1D. The strain stiffening response is clearly seen in this graph. To be consistent with our prior work(8, 10), we defined the differential shear modulus,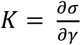, at maximum strain rate as shown in Figure 1D, where *σ* and *γ* are the measured shear stress and applied shear strain respectively. We then plot the differential shear modulus as a function of the strain in Fig. 1E for all three ECMs. A clear strain stiffening behavior is shown for all three ECMs (Fig. 1E), where the slope of differential shear modulus, *K* versus shear strain increases with the strain. The three important parameters that characterize the mechanical properties of the cogel are extracted from Fig. 1E; they are: i) initial differential shear modulus, *K*_0_, which is the differential shear modulus at zero strain; ii) strain-stiffening onset, *γ*_*c*_, the strain at which shear modulus increases by 10% from its initial shear modulus; iii) strain-stiffening exponent, *m*, the slope of the *log*(*σ*)-*log*(*γ*) plot beyond strain onset. This characterizes the strain-stiffening propensity of the gel as the fibers become aligned. We note that these are the three parameters that we will input to the fibrous material model later when calculating cell traction force. In Fig. 1E, and also Table S1, we see that the addition of HA lowers the stiffness of the gel (lower *K*_0_), and decreases the propensity of being aligned (higher *γ*_*c*_, and smaller *m*). To study the effect of HA on the viscoelasticity of the gel, sinusoidal fits on the oscillatory strain and stress response curve were performed (Fig. S2A). Viscoelasticity is defined by the loss factor at 0.1Hz. We found that viscoelasticity increases with i) collagen concentration; ii) presence of HA; and iii) the molecular weight of HA (Fig. S2B).

**Figure 1.**
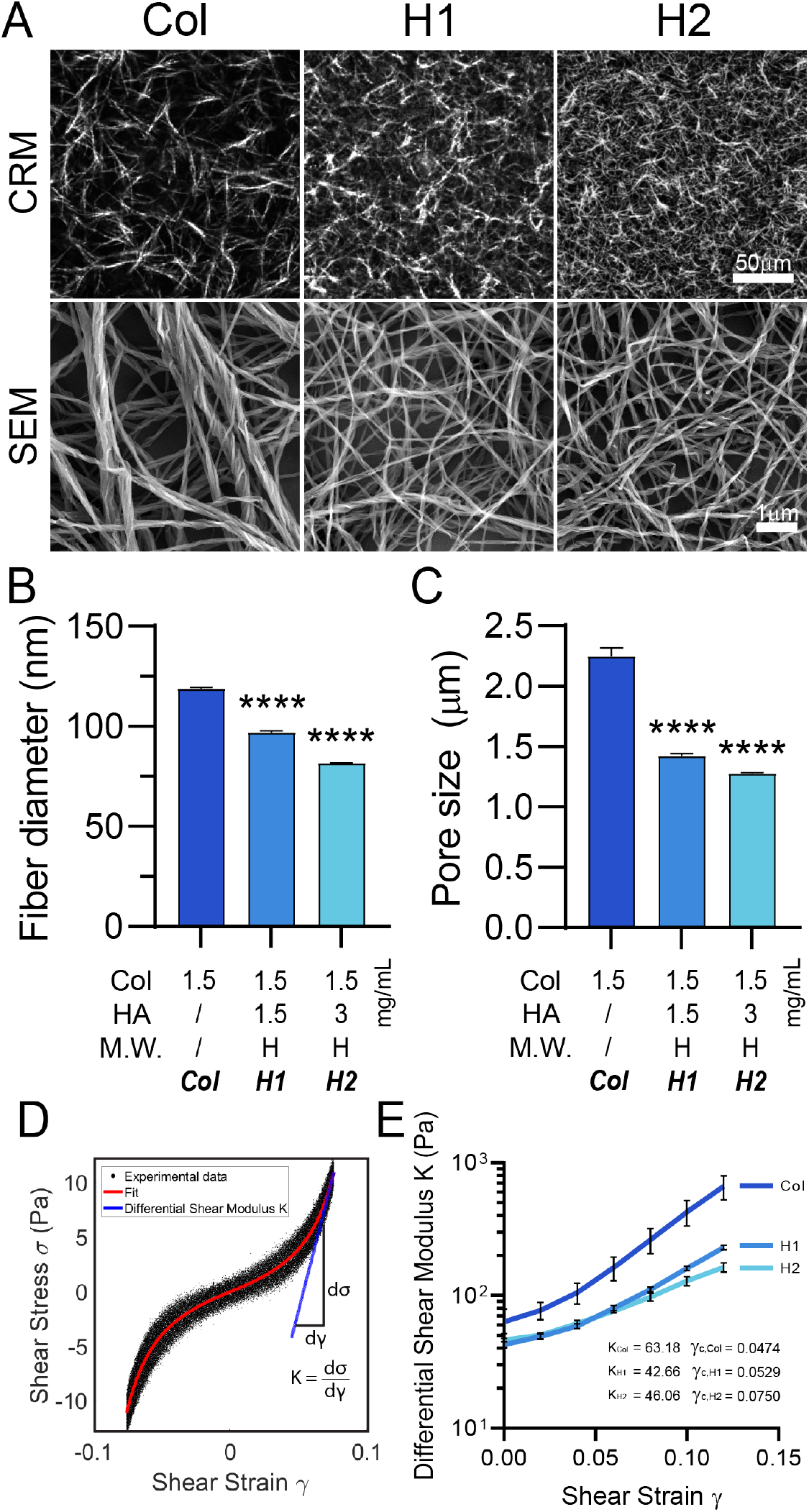
Addition of hyaluronic acid modulates the architecture and bulk mechanical properties of the collagen-HA cogels. **A**. Reflective confocal microscopy (CRM) (top row) and scanning electron microscopy (SEM) (bottom row) images of collagen fibers of matrices with Col: 1.5mg/mL collagen I only (left); H1: 1.5mg/mL collagen I and 1.5mg/mL HMA HA cogel (middle); H2: 1.5mg/mL collagen I and 3mg/mL HMW-HA cogel (right). **B-C**. Quantification of matrix architecture. HA reduces the pore size (B) and fiber diameter (C) of the collagen matrix using a porosimetry analysis (***: p<0.001; ****: p<0.0001, One-way ANOVA). **D**. Stress strain curves obtained using a parallel-plate oscillatory shear rheometer configured at 0.1Hz. Differential shear modulus is defined as, *K* = *dσ*/*dγ*, at maximum strain amplitude (left). **E**. Differential shear modulus as a function of shear strain. Here, the initial shear modulus, *K*_0_, is defined as the differential modulus at zero strain, and stiffening onset strain, *γ*_*c*_, is defined as the strain at which the differential shear modulus increases from its initial modulus, *K*_0_, by 10%.

As collagen polymerization consists of the fibril assembly and cross-linking phases(39), it is likely that the presence of HA sterically inhibits the molecular assembly of collagen fibers from fibrils by reducing the number or strength of collagen fibril-fibril interactions, leading to a decrease in fiber size(40). Indeed, it has been reported that the presence of HA in collagen matrices increases collagen fibril self-assembly rate, leading to thinner fibers as assembly rate is found to be negatively correlated with fiber size(41, 42). These thinner fibers may therefore become more susceptible to shear and compressive loads(43). According to the semiflexible polymer theory, thinner polymer fibers are likely to have lower persistence lengths, which favors bending within the network and results in smaller pore sizes(44-46). This is consistent with the architecture of the two types of gels seen in Fig. 1C, in which the H2 gel has smaller pore size and is more homogenous, which makes it less likely to be aligned. Therefore, a higher strain is required to align fibers and subsequently stiffen the gel, as indicated by a higher *γ*_*c*_.

We note that the decrease of the initial differential shear modulus in the presence of HA in Fig. 1 is consistent with previous report from Bouhrira *et al*(47) in that the addition of HA lowered the shear modulus of a collagen (0.5mg/mL) and fibrin (1.0mg/mL) cogel about 25% when all ECMs were prepared under and stayed at buffered conditions. This is consistent with the notion that HA is a filler material such that it lowers the shear force under same strain rate. Interestingly, it has been reported that HA can increase bulk and shear modulus of HA-collagen cogels when the gel is pre-swollen in a hypotonic solution (pure water) for 24hrs(18). In our experiments, however, we observed softening of the matrix under buffered condition (which is physiologically realistic) at comparable collagen and HA concentrations when no pre-swelling was performed (Table S1).

### HA impedes cell traction force generation and force transmission distance, revealed by a 3D traction force microscopy technique

A 3D traction force microscopy technique previously developed in our labs(8, 10, 18) was used to measure the single cell traction force (Fig. 2). Briefly, MDA-MB-231 cells were embedded in ECMs that were pre-mixed with fluorescent beads covalently bonded to collagen fibers(8). After overnight incubation, the cells and beads were imaged twice, before and after the addition of cytochalasin D. Upon drug treatment, actin filaments within the cell depolymerized, and the cell body relaxed. The subsequent matrix deformation was obtained by tracking the bead displacements using the two fluorescent bead images taken. The matrix deformation is shown in Fig. 2A1 and 2B1 - each arrow is a bead displacement with its length and color coded for magnitude. It is clear that the cell in the collagen only gel generated a larger deformation of matrix around the cell, and this deformation propagated further away from the cell in contrast to that in H2 (Fig. 2A-B).

**Figure 2.**
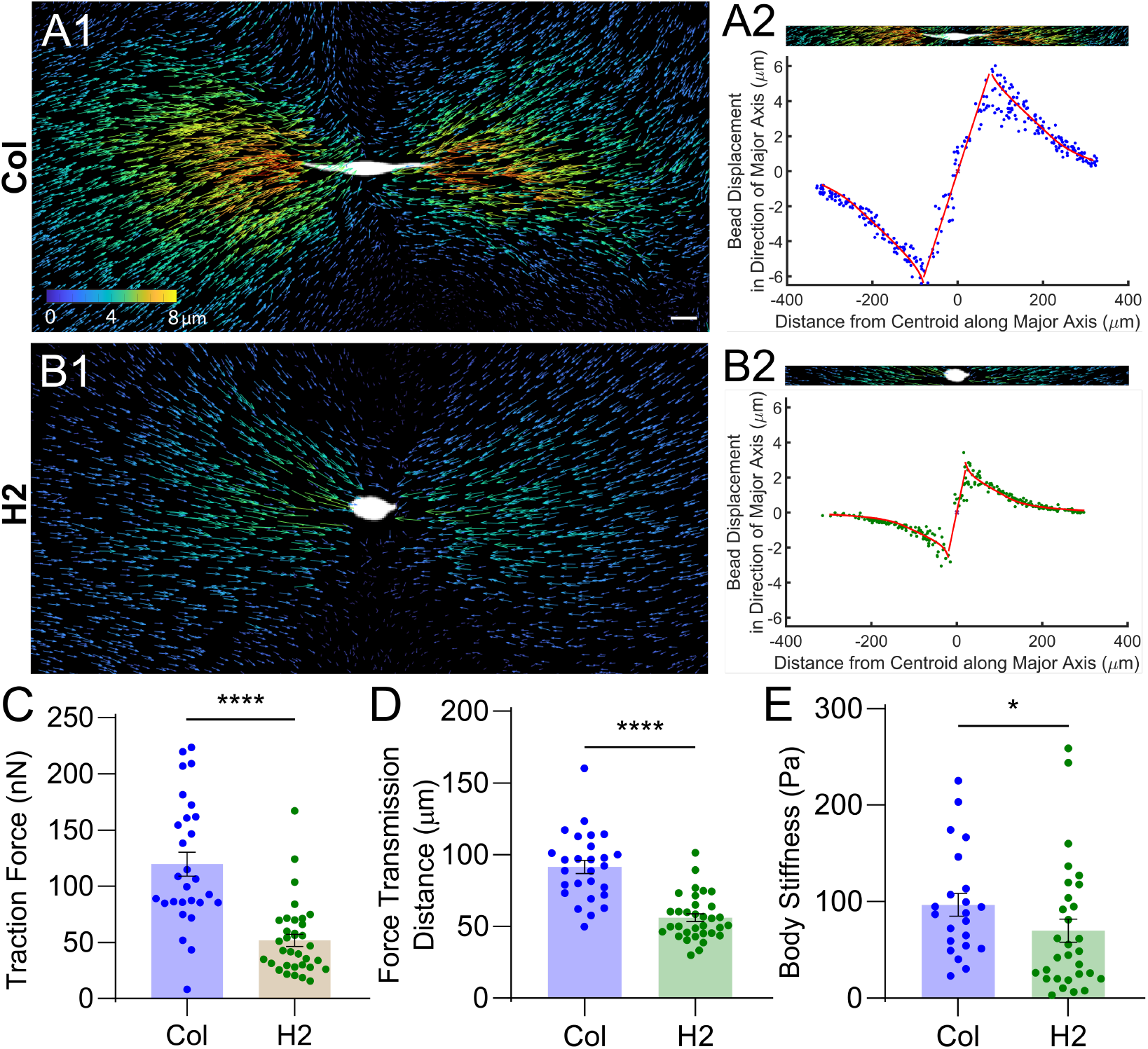
HA modulates the traction force generation of single cells. **A, B**. The ECM deformation field around a single MDA-MB-231 cell (white) embedded in collagen only gel (A) and H2 cogel (B). A1, B1: Composite images of a cell and the discrete displacements of fluorescent beads computed using the two images taken before and after cell traction force is relaxed by cytochalasin D. Each arrow represents the displacement of one bead, and the magnitude is coded by the arrow size and color for visualization. The arrows are enlarged four times their true sizes for visualization purposes. Scale bar = 20μm. A2, B2: Bead displacements along a cell’s major axis. The origin, marked by an ‘X’, represents the centroid position of the cell, and the bead displacement is computed within a 15-μm-radius cylindrical region. Dots are experimental data. Solid lines are fits to the experimental data (far field) using the constitutive fiber network model developed by Wang et al(10) (red line). The fitted parameters are used to compute the cell traction force, and force transmission distance. Near-field data is fitted with a linear function, where the cell body strain is obtained. **C**. Traction forces of cells in collagen only and H2 cogel obtained from the fits in (B). Cells in collagen only gel exert more force on the matrix, compared to cells in H2. **D**. Force transmission distance is significantly decreased in the presence of HA. The force transmission distance of a cell is defined as the distance at which the bead displacement is reduced by half from its peak. **E**. Cell body stiffness estimated by dividing cell traction force and cell body strain. Cell body strain is obtained by the slope of the near-field data in (B). Cells are stiffer in Col than in H2. All statistical tests were Mann-Whitney tests (*: p<0.05; ****: p<0.0001).

To calculate the cell generated traction force, we used the bead displacement data together with the material properties, initial shear modulus *K*_0_, onset for strain stiffening *γ*_*C*_, and the slope of strain stiffening *m* (also see Fig. 1), in conjunction with a constitutive fibrous network model previously developed in our labs(8, 10, 18). The basic idea of the traction force microscopy computation is to iteratively calculate the matrix deformation using the material properties and an assumed cell traction force until the deformation field matches the measured displacements using an optimization method. We note that it is important here that we allow matrix properties to vary slightly around the measured value in this optimization process to reflect the matrix remodeling process. The fitted displacement results are shown as solid lines (red) in Fig. 2A2 and B2.

Using the fitted parameters, we compute the cell traction force and the traction force transmission distance. The latter is defined as the distance from centroid of the cell to the half decay point of the fitted bead displacement curve illustrated in Fig. 2A2 and B2. We found that, in the presence of HA, both traction force and force transmission distance decreased in contrast to the case with collagen only gel (Fig. 2C-D). Using the cell body strain measurements (the slope of near field displacements) and cell traction force, body stiffness was estimated. We found that the cell body became softer in the H2 cogel in contrast to collagen only gel (Fig. 2E). In parallel to this, we have also demonstrated a significant decrease in traction force exerted by tumor spheroids in the presence of HA (Fig. S3).

Experimental results demonstrated that HA in a collagen gel had a profound impact on cell force generation as well as cell body mechanics. In collagen-HA cogels, in contrast to collagen only gels, cells were found to generate less traction force, the force was transmitted through a shorter distance and the cell body was softer. This is consistent with previous theoretical calculations that force transmission distance decreased in the presence of HA(18). The traction force decrease in soft gel (H2) is consistent with previous results from our labs that a stiff gel promoted cell traction force generation in 3D(8). It is likely that the cell contractility due to actomyosin was modulated in the presence of HA. It has been reported that cells in a soft ECM had weaker cytoskeletal tension due to weaker RhoA/ROCK signaling(48). As matrix stiffness decreases and fibers become thinner, cells can no longer form large, stable adhesions to develop the stress fiber network to acquire cell body stiffness and exert forces to pull the matrix(49, 50).

### CD44 is required for traction force generation in an HA-rich environment

Cell surface adhesion receptors mediate cell traction force generation by linking the cytoskeletal molecules with the fiber network in the ECMs (Fig. 3A). In collagen only gel, cells can use the surface receptor β1-integrin to link F-actin within the cell to collagen fibers within the ECM for force generation. While it has been reported that the cell surface adhesion receptor CD44 can mechanically link actin filaments with HA molecules, it is unclear whether CD44 is involved in traction force generation and transmission. Moreover, CD44 has been found to be overexpressed in triple-negative breast cancer(51). Using MDA-MB-231, a cell line known to express high levels of CD44(52), we asked the question of whether/how CD44 is involved, together with β1-integrin in cell traction force generation, thereby potentiating tumor invasion.

**Figure 3.**
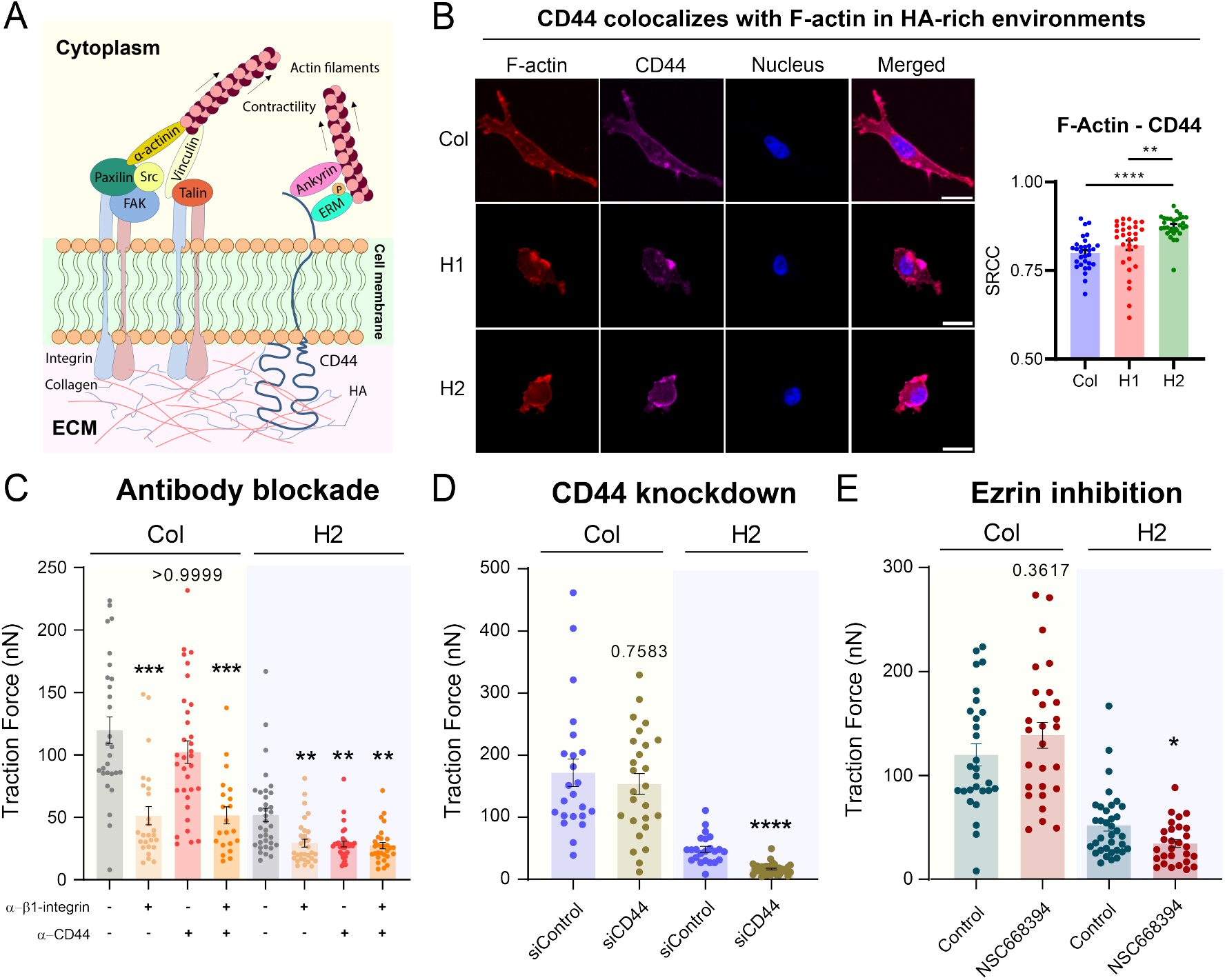
CD44 is required for traction force generation in an HA-rich environment. **A**. An illustration depicting the engagement of both CD44 and *β*1-integrin to actin and their adhesion to ECM components. **B**. Colocalization of F-actin and CD44. Maximum intensity Z-projections of fluorescence images of F-actin, CD44, and the nucleus of MDA-MB-231 cells embedded in the three gels (left panel). Colocalization is quantified by the Spearman’s rank correlation coefficient (SRCC) (right panel). Analyses were performed over all the Z slices of confocal images taken from individual cells. Significantly higher colocalization between F-actin and CD44 in H2 was found. Scale bar = 20 *μ* m. **C**. Cell traction force when CD44 and/or *β* 1-integrin was neutralized by antibodies in Col and H2. While cells in Col are insensitive to CD44 blockade, blocking of *β*1-integrin leads to a significant decrease in traction force. In H2, blocking either *β*1-integrin or CD44 was sufficient to reduce traction force significantly. **D**. CD44 knockdown resulted in a significant reduction in cell traction force in H2, but not in Col. **E**. Cell traction force when cells were treated with NSC668394, a small molecule inhibitor for ezrin at Thr567, which disengages ezrin-mediated actin-CD44 interactions. Cell traction force reduces significantly when treated with the ezrin inhibitor in H2, but not in Col. All statistical tests were Kruskal-Wallis tests (*: p<0.05; **: p<0.01; ***: p<0.001; ****: p<0.0001).

To establish that CD44 is capable of relaying force, we imaged the colocalization of CD44 and F-actin in MDA-MD-231 cells embedded in the three gels. In H2 cogels, there was significantly more colocalization between CD44 and F-actin, compared to cells in Col and H1 (Fig. 3B). While more colocalization was found between CD44 and F-actin, F-actin seem to reside preferentially on the periphery of cells in H2, compared to stress fibers observed in cells embedded in Col (Fig. S4).

To test whether CD44-mediated adhesion is required for force generation in addition to β1-integrin in the presence of HA, we performed cell traction force measurements of cells when either or both CD44 and β1-integrin were neutralized by monoclonal antibodies. To ensure fair comparison, control groups are treated with IgG-1 to account for non-specificity. In Col gel, traction force was significantly attenuated when β1-integrin was blocked, regardless of CD44 blockade (Fig. 3C), suggesting that β1-integrin is the primary mechanical link for cell traction force generation in collagen only gel. In H2 gel, blocking either β1-integrin or CD44 resulted in significant traction force reduction (Fig. 3C). Interestingly, traction force did not reduce further when both adhesion molecules were blocked (Fig. 3C). We then validated the antibody blockade experiments by knocking down (KD) CD44 using a pool of three siRNAs (Fig. 3D) and performed TFM experiments in Col and H2. We found that CD44 knockdown results in significant reduction of traction force in H2, but not in Col (Fig. 3F). This is consistent with our flow cytometry analysis on phospho-myosin light chain (p-MLC), the downstream effector of the Rho-ROCK pathway for actomyosin contractility, in which we found a downregulation of p-MLC in H2 but not in Col when CD44 is knocked down (Fig. S5A-B). We also assessed whether ezrin-mediated actin engagement to CD44 may serve as a mechanical link for force generation by inhibiting ezrin phosphorylation at Thr567, a known actin-binding site(53). When treated with NSC668394, a small molecule inhibitor for p-ezrin(Thr567), traction force reduces significantly in H2, but not Col (Fig. 3E).

Taken together, Figure 3 shows (i) cell traction force generation via CD44 is as crucial as that via β1-integrin in HA; (ii) HA inhibits cell traction force generation. This is consistent with prior work that stiffer environments promote traction force generation(8). A FRET experiment by Yang et al. showed that HMW HA can induce aggregation of CD44(54), which may explain why cells can still exert forces via CD44 even though forces acting on each CD44 may be weak. (iii) the forces generated through the CD44 in contrast to β1-integrin are likely to be weaker (Fig. 3C) as HA fibers are much softer than collagen fibers(40). Previous work has indicated the possible linkage between CD44 and β1-integrin, in that they are co-enriched at adhesion sites via lipid raft coalescence, and the activation of CD44 can induce upregulation of β1-integrin(55-57). Nonetheless, we are not ruling out the possibility that HA may affect traction force generation through other transmembrane collagen receptors that are tethered with the actin cytoskeleton, such as discoidin domain receptor 1 (DDR1)(58, 59), which may also contribute to the residual traction we observed when β1-integrin and/or CD44 was modulated. It is well-known that receptor for hyaluronan-mediated motility (RHAMM) also interacts with the actin cytoskeleton(60, 61). However, we suspect that traction force is less likely to be mediated through RHAMM, compared to CD44, due to its relatively lower binding affinity for HA(52).

### HA modulates cell-EM interactions via rhoA mediated contractility leading to less fiber alignment

One of the hallmarks and physical cues for tumor cell invasion during cancer metastasis is collagen fiber alignment due to matrix remodeling(62, 63). It has been shown that highly aligned collagen fibers can facilitate cell dissemination through directional migration(64, 65), and/or cell-cell communication(18, 65). While our rheological findings suggest that the presence of HA decreases collagen network pore size, we examined whether fiber alignment due to weaker cell traction force is also impaired in the presence of HA. Figure 4A shows reflective confocal images of collagen fibers surrounding an MDA-MB-231 cell embedded in collagen only (Col), and collagen-HA cogels (H1 and H2). It is evident that cells are more rounded in both H1 and H2 co-gels than in Col. Furthermore, collagen fibers are more bundled together, have longer persistence length, and aligned around the tip of cell in Col. This alignment is less pronounced in H1 and H2 cogels. To quantify the above observation, we calculated the aspect ratio of cells. In Fig. 4B1, the aspect ratios of the cells are significantly larger in Col (6.68±3.69) than H1 (2.07±0.85) and H2 (2.36±2.03). Cells are more amoeboid like in the more homogenous collagen-HA cogels, likely a result of less mechanical cross talk between cells and ECMs (Fig. 4B1). We also observed that elongated cells tend to generate higher levels of traction forces, regardless of the matrices they are embedded in (Fig. 4B2). Using a nematic tensor-based anisotropy score method(66), we quantified the extent of fiber alignment in the vicinity of cells. The anisotropic scores indicate that fibers around cells in collagen only gel are significantly more aligned compared to fibers around cells in H2 cogels (Fig. 4B3). We also observed that fiber alignment by tumor spheroids is impaired in H2 (Fig. S6). When we probed RhoA activities during traction force generation, using a previously developed Förster resonance energy transfer (FRET) imaging workflow(67), we found that the RhoA activity appeared to be more associated with matrix deformation in Col, but not in H2 (Fig. 4C). By analyzing the cross correlation between the matrix deformation (Fig. 4D1) and RhoA activity (Fig. 4D2), we found that RhoA activity in Col is correlated to the bead displacement around the cell (Fig. 4D3), suggesting that matrix deformation is caused by actomyosin contractility in Col, but not in H2. This suggests that HA limits the mechanical communication between cells through modulation of RhoA-mediated contractility.

**Figure 4.**
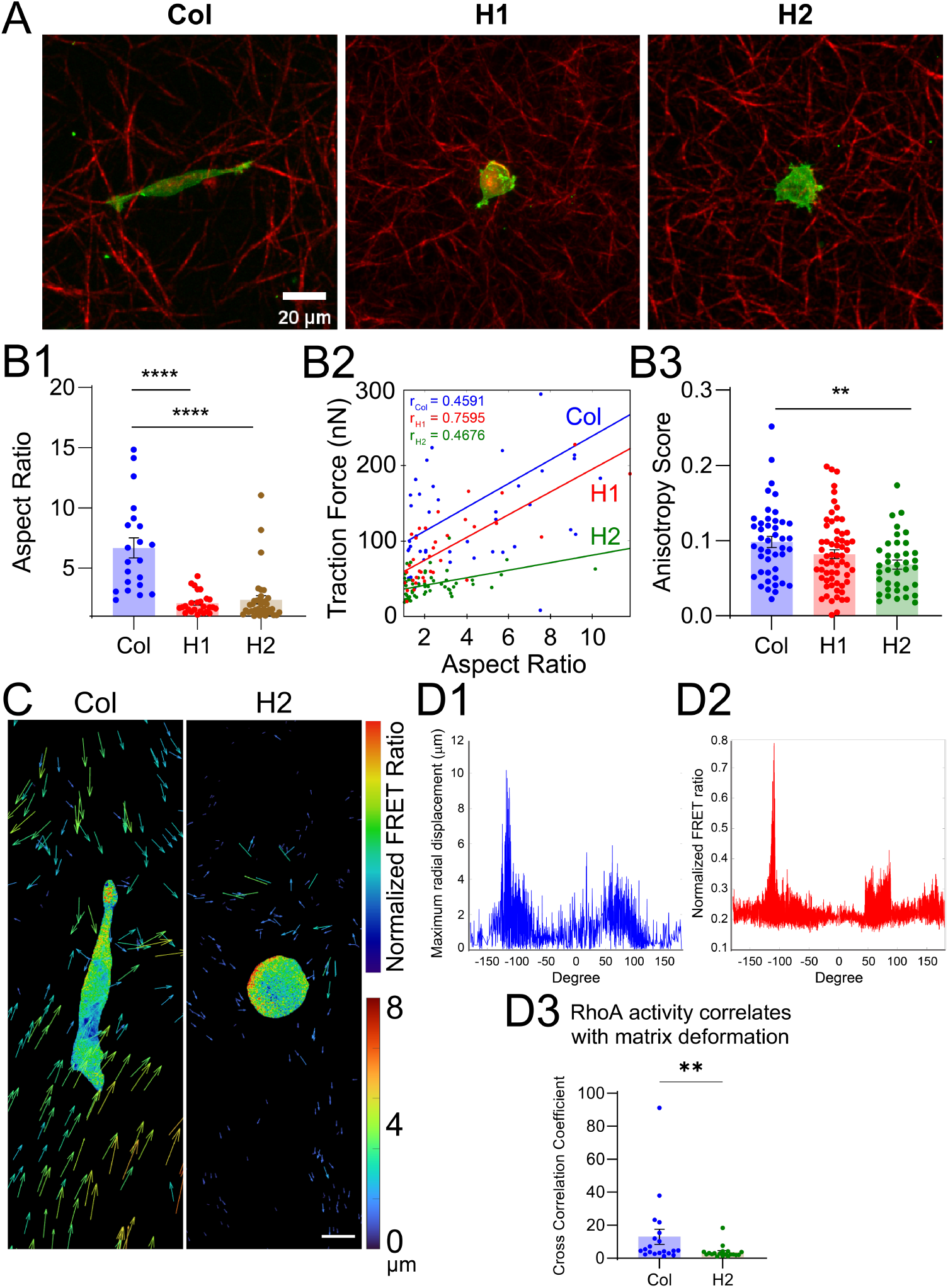
Cells in collagen are more elongated, and have polarized RhoA activity, leading to fiber alignment. **A**. An overlay image of a GFP-actin tumor cell (green) and a reflective confocal image of the collagen fibers (red) in three different gels. **B1**. Cells are more rounded in collagen-HA cogels in contrast to collagen only gel. (n=21 for Col; n=26 for H1; n=36 for H2). **B2**. Elongated cells generate higher traction forces in all matrices as shown by the Pearson correlation coefficients *r*. **B3**. Collagen fibers around the tips of the cells are significantly less aligned in H2 cogel than in pure collagen gel. Quantification of fiber alignment is done using the ImageJ plugin, FibrilTool. 15 *μ* m-radius cylindrical regions at the two extremities of cells are chosen for calculation of anisotropy score. (n=44 for Col; n=62 for H1; n=38 for H2). All statistical tests were one-way ANOVA tests (**: p<0.01; ****: p<0.0001). **C**. RhoA activity and matrix deformation revealed by a TFM-FRET technique. Scale bar = 10 *μ*m. **D1**. Radial profile of maximum bead displacement cells. **D2**. Radial profile of RhoA activity measured by FRET ratio. **D3**. Cross correlation analyses between bead displacement and FRET ratio profiles show that RhoA activity in Col is correlated to the deformation around the cell, suggesting that matrix deformation is caused by actomyosin contractility in Col. Statistical test was Mann-Whitney test (**: p<0.01).

Here, we have quantitatively demonstrated *in vitro* that the presence of HA in a collagen network limits fiber alignment in single cells, which may be correlated with the modulation of RhoA-mediated contractility. This is consistent with previous theoretical predictions. Theoretical work from the Shenoy lab showed that the presence of GAG decreased the propensity of a swollen collagen network to be aligned(18). Numerical simulations and microparticle experiments performed by Hatami-Marbini et al. have suggested that composite collagen networks consisting of high concentrations of HA are less favorable to fiber reorientation and allow shorter distance propagation of matrix deformation(68).

### HA attenuates tumor invasiveness, and breast tumor invasion is CD44-dependent, as revealed in both 3D single cell and tumor spheroid invasion assays

Lastly, we investigated how tumor cells migrate and invade within an HA-rich matrix in contrast to a collagen only matrix using a single cell and tumor spheroid migration assay. We embedded single MDA-MB-231 cells into the matrices and tracked cells with an in-house MATLAB algorithm to compute migration speed and mean squared displacement (Fig. 5A-B). We found that the presence of HA significantly reduced single cell motility (Fig. 5A-B). To model invasion from a primary tumor mass, we embedded tumor spheroids into the matrices to study spheroid invasion behavior (Fig. 5C-D). We quantified invasiveness by computing the percentage of cells escaping from the spheroids (Fig. 5D1) and the expansion of spheroids (Fig. 5D2). We found that tumor spheroids were less invasive in H2 than those in Col (Fig. 5C-D). Knowing that CD44 is required in generating force in a HMW HA-rich environment, and that tumor invasion requires force, we questioned whether tumor invasiveness can be modulated by CD44. We therefore pre-incubated tumor spheroids with NSC668394 (the ezrin inhibitor) and *α*-CD44 respectively prior to invasion assays, and followed tumor invasion for 16hrs (Fig. 5E). In both treatments, we observed less cells invading out of the tumor cores in H2, but spheroids in Col were unresponsive to treatment (Fig. 5F). We then quantified invasion by the expansion of spheroids, and found that spheroid expansion was significantly reduced by ezrin inhibition and CD44 blockade (Fig. 5F).

**Figure 5.**
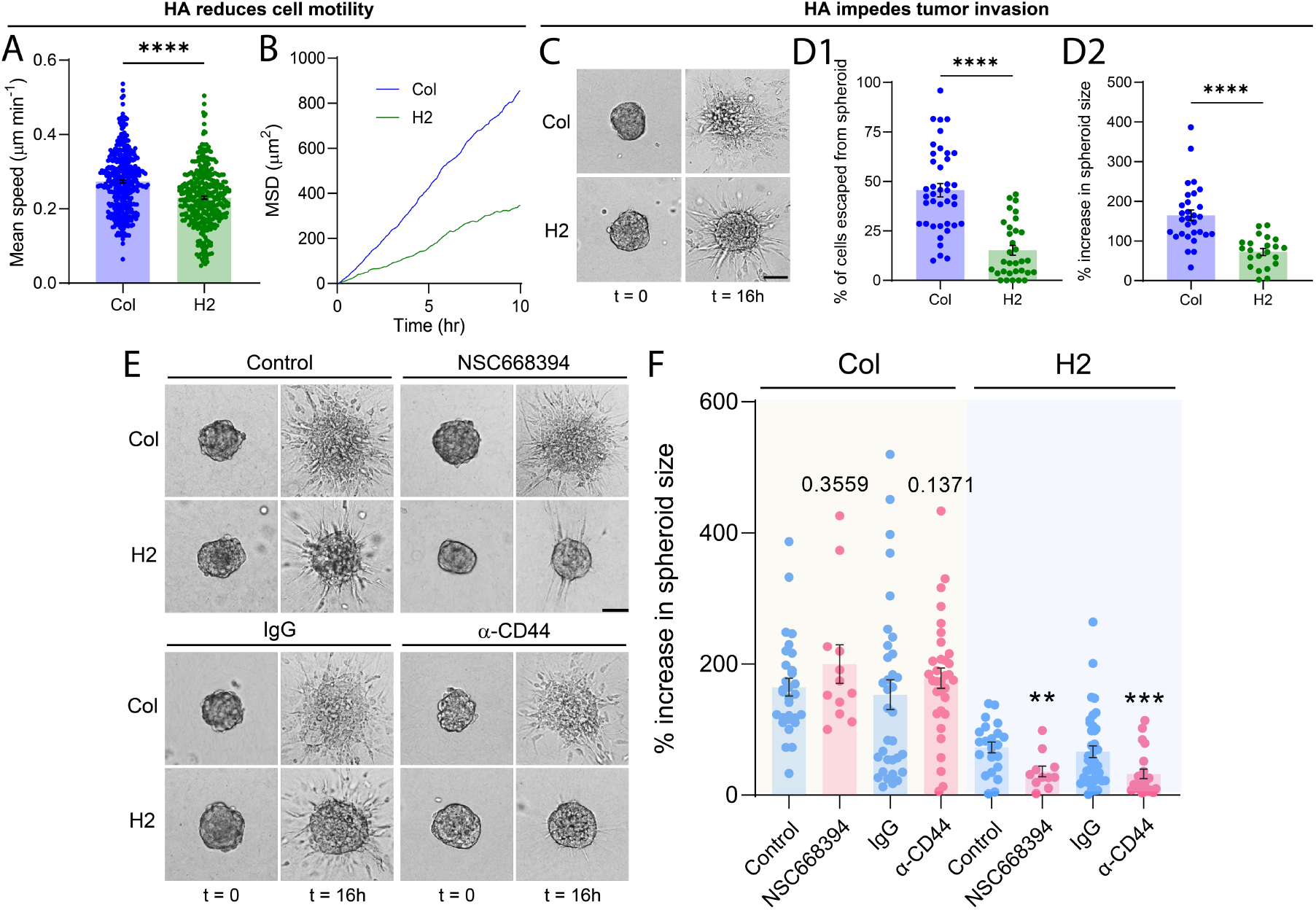
HA impedes cell migration, and tumor invasion in an HA-rich environment is CD44-dependent. **A-B**. Quantification of cell motility in MDA-MB-231 single cells using mean speed (A) and mean squared displacement (MSD) (B). **C**. Spheroid invasion assay using MDA-MB-231 spheroids. Scale bar = 50*μ*m. **D**. Quantification of tumor invasiveness. We counted the percentage of cells escaped from the initial spheroid boundary (D1) and the expansion of spheroids (D2). Tumor spheroids are less invasive in the presence of HMW HA. **E**. Tumor invasion when ezrin phosphorylation was inhibited (top) and when CD44 was blocked (bottom). Scale bar = 50*μ*m. **F**. Quantification of tumor invasiveness using the percentage increase in spheroid size. Both ezrin inhibition and CD44 blockade had no effect on tumor invasion in Col, while tumor invasion in H2 was reduced significantly by CD44 blockade. All statistical tests were one-way ANOVA tests (**: p<0.01; ***: p<0.001; ****: p<0.0001).

Here, we have demonstrated that HMW HA impaired the tumor cells’ ability to invade within ECMs in both single cell and tumor spheroid assays. This is consistent with our ECM characterization and traction force measurements. We showed that cells generated less traction force in HA-rich gel, thus less likely to align collagen. This is also reflected in the spheroid invasion assay in that ECMs were observed to be more aligned radially in Col in contrast to H2, facilitating invasion (Fig. S6). Our work has also revealed that tumor invasion in an HA-rich environment can be reduced by interfering with CD44. This can be explained by the fact that cells can generate traction force through CD44, which may enable cell motility and thus the breakout from tumor spheroids.

It is well-documented that cells can produce HA with different molecular weights(69), and it has been reported that modulus increases monotonically with the molecular weight of HA, and helical structures have been observed in HMW (>1MDa) HA gels(40). Therefore, it is possible that force generation through CD44 depends on the molecular weight of HA, thereby influencing tumor invasion. We note that the HA we used here is HMW HA, and there have been reports that HMW HA and LMW HA inhibits/promotes single cell migration(69). We observed that tumor invasion is promoted in Col-LMW HA cogels (L2) compared to Col and H2 (Fig. S7 and Fig. 5C-D). This is likely due to the fact that cells in L2 generates much higher levels of traction force, compared to Col and H2 (Fig. S8), and L2 matrix has a higher propensity to strain stiffen 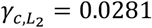 vs *γ*_*c,Col*_ = 0.0474 and 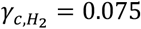 (Table S1), which are mechanically favorable to cell migration. However, the question of whether cell migration is mechanically driven by CD44 in LMW-HA-rich environment remains unanswered. Our data shows that CD44 modulation by antibody blockade and ezrin inhibition had minimal effect on tumor invasion (Fig. S7), and CD44 knockdown, but not blockade, significantly reduces traction force generation (Fig. S8). These findings suggest that the mechanical linkage between CD44 and LMW HA has little effect on tumor invasion, while it is possible that CD44 is involved in other mechanisms that promotes cell migration. For example, the trans-activation of *α*_5,_*β*_1-_ -integrin by CD44 and its subsequent phosphorylation of paxillin as demonstrated by McFarlane *et al*(55), which stabilizes focal adhesions for effective migration.

## Materials and Methods

### Cells, spheroids, and matrices

#### Cells

Triple-negative breast tumor cells (MDA-MB-231) (ATCC) were used in all experiments. Cells were maintained at 37°C and 100% humidity in a 5% CO_2_ incubator and were cultured in T75 flasks (10062-860, Corning) using Dulbecco’s Modified Eagle’s Medium (DMEM) (12430054, Gibco) with 4.5g/L glucose, L-glutamine, and N-2-hydroxyethylpiperazine-N’-2-ethanesulfonic acid (HEPES) supplemented with 10% fetal bovine serum (FBS) (26140079, Gibco), 1% penicillin/streptomycin (15140122, Gibco), and 1mM sodium pyruvate (11360070, Gibco). Cells were passaged at 70-90% confluency. Cells with 13-20 passages were used for the experiments. For cell tracking purposes, MDA-MB-231 cells were labelled by the Lammerding lab using the lentiviral construct pCDH-H2B-mScarlet(70). GFP-actin-labeled MDA-MB-231 cells were a generous gift from the Kamm lab at MIT. MDA-MB-231 cells with the RhoA FRET biosensor was previously developed in the Hodgson lab and validated(67). Briefly, the biosensor consists of a Rho binding domain (RBD) from Rhotekin, enhanced cyan fluorescent protein (ECFP), an unstructured linker, Citrine yellow fluorescent protein (YFP), and the full length RhoA GTPase, in a single-chain construction(71). Biosensor expression was regulated by tetracycline-inducible expression system (tet-OFF: Clontech) and stably traduced using a retrovirus(72). The cells were selected for stable genomic integration using 10 μg/mL puromycin dihydrochloride (A1113802, Gibco) and 1 mg/mL G418 (10131035, Gibco) in culture media, together with 1 μg/mL doxycycline hyclate (D9891, Sigma) to repress the biosensor expression under normal culture. The culture media for this cell line is Dulbecco’s Modified Eagle’s Medium (DMEM) (15-013-CV, Corning) with 4.5 g/L glucose, sodium pyruvate supplemented with 1:100 GlutaMAX (35050061, Gibco), 10% fetal bovine serum (FBS) (S11150, Atlanta Biologicals), 1% penicillin/streptomycin (15140122, Gibco), and 1 mg/mL G418 (10131027, Gibco). To induce expression of the biosensor, 48hrs prior to experiments, cells were trypsinized and doxycycline was removed to allow expression in doxycycline-free complete medium.

#### Spheroids

Tumor spheroids with uniform size were generated using a microfabricated microwell array platform previously developed in our lab(73). Briefly, 36-by-36 microwell arrays were patterned in thin agarose 1cm-by-1cm squares. These squares were placed in 12-well plates, one square for each well. Within each well of the 12-well plate, 2×10^6^ cells (4:1 ratio of non-labeled MDA-MB-231:mScarlet-histone H2B-labeled MDA-MB-231 suspended in 2.5mL complete medium were introduced. This spheroid growth medium is different from the culture medium used in single cells and supports spheroid formation. It was composed of DMEM/F-12 medium (11320033, Gibco), 5% donor horse serum (26140079, Gibco), 20 ng/mL human EGF (PHG0311, Gibco), 0.5 µg/mL hydrocortisone (H0888–1G, Sigma-Aldrich), 0.1 µg/mL cholera toxin (C8052-.5MG, Sigma-Aldrich), 10 µg/mL insulin (I0516-5 ML, Sigma-Aldrich), and 1% penicillin/streptomycin (Gibco). Cells were allowed to settle and form spheroids in microwells measuring 200μm by diameter and 220μm by height for 5 days. To maintain a sufficient level of nutrients, medium was changed on the 3rd day after seeding. Spheroids with sizes ∼100μm were generally formed in these microwells after 5 days.

#### 3D cell and spheroid culture

To render a 3D environment for cells, rat tail type I collagen (354249, Corning) in 0.1% acetic acid was used. On ice, 31.65μL collagen stock (9.48mg/mL), 20μL 10X M199 medium (M0650, Sigma-Aldrich), and a pre-titrated volume of 1M NaOH (79724, Sigma-Aldrich) were mixed with the cell suspension in complete medium to reach a final volume of 200uL and a pH of 7.4. The final concentration of collagen in all gels was 1.5mg/mL. To prepare HA stock solution for cogelation with collagen, sodium hyaluronate powder (HA15M-1 and HA10K-1, Lifecore Biomedical) was reconstituted in phosphate buffered saline (10010023, Gibco) to reach a final concentration of 10mg/mL. The HA stock solution was added to the collagen mixture for cogelation at final concentrations of 1.5mg/mL and 3.0mg/mL to yield 1:1 and 1:2 collagen: HA ratios respectively. Matrices with cells and spheroids were prepared using methods described in the methods section. For TFM experiments, single cells were seeded into the ECM at a cell density of 50 cells/μL. More details are provided in SI Materials and Methods for spheroid harvesting, cell and spheroid embedding, and neutralization of adhesion molecules. Collagen matrices were placed in an imaging device described in *SI Materials and Methods* and Fig. S9.

### Mechanical characterization of ECMs

#### Characterization of bulk mechanical property of ECMs

Discovery HR-3 Shear Rheometer (Texas Instruments, USA) was used for mechanical characterization of gels with a 25-mm parallel-plate geometry and 500μm trim gap (Fig. S1). Briefly, two 25mm No. 1.5 cover glasses (630-2213, VWR) were surface-treated with glutaraldehyde and bonded to each plate of the rheometer with double-sided tape (666, 3M). The rheometer was then zeroed, calibrated, pre-chilled to 4°C for *in situ* gel polymerization. 270μL ECM solution was pipetted onto the parallel plate with a trim gap of 500μm. The solution was then carefully sealed with mineral oil (330779-1L, Sigma-Aldrich) to prevent evaporation throughout the experiment. It is important to note that any movement on the ECM setup would cause pre-alignment of collagen, leading to inaccurate measurements. We also made sure that the ramping temperature for ECM polymerization is the same as that of real experiments (Fig. S10).

Gel polymerization progress was then monitored by imposing three cycles of 0.5% strain every 5 min to measure the shear storage modulus, G’, as a function of polymerization time. Strain-stiffening tests were carried out after the shear storage modulus reached its plateau. The matrices were subjected to 10 oscillations at 0.1Hz at increasing shear strain amplitudes from 2 to 12% strain in 2% increments and from 12 to 40% in 4% increments. Note that the strain amplitudes here are the maximum strain amplitudes of the circular parallel plate during shear (i.e. at the edge of the plate). To define the stiffnesses of matrices, the differential shear modulus 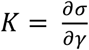 at maximum strain amplitude of each strain level was used to estimate the initial shear modulus *K*_0_ at which the strain was zero. To characterize strain-stiffening behavior, we used the stiffening onset strain, *γ*_*c*_, which is the strain at which the shear modulus increases from its initial modulus by 10%(8). As such, a lower *γ*_*c*_ would indicate a stronger strain-stiffening behavior.

#### Imaging of collagen matrices

The ECMs were imaged using a reflective confocal imaging modality with a 640nm laser, a C-Apochromat 40x/1.20 W Corr M27 objective (Zeiss) and a LSM710 Zeiss confocal microscope. Typically, each image is 212.55 × 212.55 × 100 μm in size with a voxel size 0.415 × 0.415 × 0.5μm (x, y, and z respectively). To process the images for downstream analyses, images were background subtracted in ImageJ (NIH) with a 10-pixel radius, followed by a 3D Gaussian filtering step with a sigma of one voxel. The images were then auto-thresholded using methods by Huang and Wang(74). The image stacks were further skeletonized using the Skeletonize3D plugin in ImageJ (NIH).

#### Pore size and fiber diameter estimation

Using the skeletonized binary images, size distribution of pores were estimated using a porosimetry-based analytic technique in PoreSpy, a Python toolkit developed by Gostick et al(75). The pore size distribution was then fitted with a Rayleigh function and the average pore size was defined by the mean of the fitted Rayleigh distribution. To correct for the invisible fibers due to a blind spot in confocal reflectance microscopy(76), a scaling factor of 1.82 was applied to the Rayleigh mean to yield the estimated average pore size. To estimate fiber diameter, we adopted a volumetric approach, as previously described, where fiber thickness was estimated by the total volume and length of fibers(8). First, the total length, *L*, of all fibers in a matrix was obtained using the *AnalyzeSkeleton* function. To estimate the total volume, *V*, of fibers, we assumed that the material density of collagen fibers to be close to water, 1g/cm^3^, and calculated the total volume of fibers by knowing the concentration of collagen in the matrix. By considering the geometrical shape of collagen fibers as perfect cylinders, fiber diameter, d, was then given by the mathematical expression 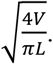.

#### Quantification of fiber alignment

Fiber alignment was analyzed within two 15-μm radius-cylinders along the long axis of a cell at two cell tips. Using a nematic tensor-based ImageJ plugin, FibrilTool(66), the anisotropy of fibers was quantified, where regions with high anisotropy scores indicate high levels of fiber alignment.

### 3D traction force microscopy (TFM)

Cell traction force was computed using the matrix deformation field generated near the cell, the bulk material properties measured from rheometer, and a constitutive fibrous nonlinear elastic model previously developed in our labs [18,73]. Two critical steps of the TFM are (i) to obtain the ECM deformation field around the cell; (ii) to translate the ECM deformation along with ECM mechanical properties into cell traction force. To facilitate matrix deformation measurement, 1μm-diameter fluorescent beads were coupled to and uniformly distributed in the cell embedded ECM. Two image stacks of the florescent beads were taken (See SI Materials and Methods) before and after the cell was relaxed by cytochalasin D. We used 10μM cytochalasin D (C8273, Sigma-Aldrich) to disrupt the actin filaments and cells were given 1hr to relax.

To extract the bead positions, we used a maximum likelihood estimator-based Lucy-Richardson deconvolution method. To compute the bead displacement, we used a topology-based particle tracking (TPT) algorithm(77). We then plotted bead displacement, s, along the major axis of the cell within a 15-μm-radius cylinder lengthwise. To translate matrix deformation into cell traction force, using the constitutive model for fibrous nonlinear elastic gel previously developed in our labs(8, 10, 18), we numerically calculated traction force such that the resulted displacements fit to the experimentally measured bead displacements using the Nelder-Mead search method implemented in MATLAB. More specifically, the input parameters are ECM deformation field and the mechanical properties (initial differential shear modulus, strain onset for strain stiffening and slope of the strain stiffening) of the ECM. The adjustable parameter is the principal stress along the long axis of the cell, and the ECM mechanical properties. The Nelder-Mead search algorithm minimizes the least square of the difference of the theoretically evaluated displacement field with that measured in experiments. The fitted parameter, the principal stress along the cell major axis is then used to calculate the traction force. Force transmission distance was defined as the axial distance between the cell centroid and the point where bead displacement decayed by half from its maxima. We note that cells often have sharp protrusions which deviate the shape from an ideal ellipsoid as represented in our estimations. Our previous work has illustrated that radius of curvature at cell tips does not have any significant effect on the range of displacement propagation and force magnitude(8).

### FRET imaging of RhoA activity

A dual channel image acquisition system was used to acquire the FRET and CFP image pair as previously described(67). In brief, the FRET and YFP channels were acquired simultaneously using the W-View Gemini beam splitter (A12801-01) for ratiometric FRET calculation. To generate FRET images, images from the CFP/FRET image channels were aligned, flatfield-corrected and background subtracted. Cells were segmented using a convolutional neural network package, U-Net, developed in the University of Freiburg(78). A previously trained model was used to segment cells[75]. The ratiometric FRET (*rFRET*) image was calculated by the following equation: 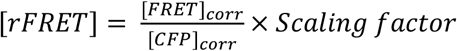 Where [*FRET*]_*corr*_ and [*CFP*]_*corr*_ are the corrected images of FRET and CFP channels respectively. The scaling factor was used to bring images to visualization under a 16-bit domain. The choice of scaling factor did not affect our final calculation as long as it was consistent for both channels.

### Cross correlation analysis between matrix deformation and RhoA activity

To correlate matrix deformation and RhoA activity of the cell, we plotted the maximum bead displacement around a cell and the average FRET ratio (I.e. RhoA activity) in all radial directions, spanning from the cell centroid. Cross correlation coefficient was then used to quantify the similarity between the two spatial profiles, thereby inferring the correlation between matrix deformation and actomyosin contractility.

### Flow cytometry analysis of p-MLC

Collagen matrices containing cells were incubated with 1 mg/mL collagenase in the presence of 1x Halt™ phosphatase inhibitor cocktail (78420, Thermo Scientific) at 37°C for 1hr. Cells were then centrifuged in a staining buffer with 2% FBS (26140079, Gibco) in PBS, followed by a 30-min incubation at room temperature in staining buffer containing 1x LIVE/DEAD fixable viability dye (L34990, Invitrogen) and 1x Halt™ phosphatase inhibitor cocktail (78420, Thermo Scientific). Cells were then washed and centrifuged, and fixed in 200uL 1.5% formaldehyde for 10mins at room temperature. After fixation, cells were centrifuged and washed in staining buffer, followed by permeabilization using 200uL ice-cold 99.8% methanol (322415, Sigma) for 15mins on ice. Cells were then centrifuged and washed twice in staining buffer. Cells were incubated with 2.5% normal goat serum (ab7481) for 10mins on ice to block non-specific binding. To stain for p-MLC, cells were incubated with 1:100 anti-p-MLC2 (Ser19) rabbit polyclonal antibody (3671, Cell Signaling Technology) for 15mins on ice, and 1:1000 goat anti-rabbit IgG H&L conjugated with Alexa Fluor 488 (ab150077) for 30mins at room temperature, with two washing steps in between. All samples were resuspended in staining buffer and kept on ice before analysis. All flow cytometry analyses were performed on the Attune NxT flow cytometer (ThermoFisher, USA). ∼20,000 events were recorded for each sample. Unstained, single stained, and Y-27632 controls were used to determine the compensation matrix and negative gates for data analysis in FlowJo (Fig. S11B).

### Invasion assay

Single cell invasion. mScarlet-histone H2B-labeled MDA-MB-231 cells were embedded in the ECMs at a cell density of 400 cells/μL in complete culture medium. Images were acquired with a 10x magnification objective lens (NA = 0.4; UPlanSApo, Olympus America) installed on an inverted epifluorescence microscope (IX81, Olympus America) and a 16-bit sCMOS camera (ORCA Flash 4.0 V3, Hamamatsu Photonics). Brightfield and fluorescent images were captured every 5 mins for over 10 hrs. Fluorescence images of cell nuclei were then thresholded by Otsu’s method and tracked using an in-house centroid tracking algorithm in MATLAB. Mean speed of a cell was defined by the total distance travelled divided by the total time. Mean squared displacement (MSD) was defined by the following formula: 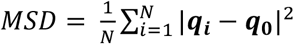, where *N* is the total number of cells, vector ***q***_***i***_ is the position of the *i*^th^ cell at time *t*, vector ***q***_**0**_ is the initial position of the *i*^th^ cell.

#### Tumor spheroid invasion

Spheroids were generated and embedded in matrices as previously described(73) (Also see *SI Materials and Methods*), followed by imaging on a climate-controlled spinning disk confocal microscope (Andor/Olympus) with a 10x objective lens (NA = 0.4; USPLSAPO10X2, Olympus America). Image stacks measuring 561.1 × 561.1 × 100 μm with a voxel size of 1.0959 × 1.0959 × 1μm were taken with a 605 nm laser every 14 mins for over 16 hrs. To quantify invasion, we computed the percentage of cells invaded out of the initial spheroid boundary after 16 hrs.

## Acknowledgments

We would like to thank Prof. Jan Lammerding for insightful discussions and advice throughout this project, Dr. Jeremy Keys for his help in transducing cells with fluorescent labels, Dr. Matthew Hall for his help in particle tracking, Prof. Jeff Gostick for his help on setting up the pore size analysis algorithm, Dr. Joanna Dela Cruz for her help on imaging, and staff at the Cornell University BRC Flow Cytometry Core Facility (RRID:SCR_021740) for their help on flow cytometry analysis. The experimental part of the work was supported by the following NIH grants: R01CA221346 (to M.W.&J.E.S), S10RR025502, S10OD010605, S10OD018516 (to Cornell BRC), R35GM136226 (to L.H.). The theoretical part of the work was support by the following NCI grants: R01CA232256, U54CA261694 (to V.B.S).

