## Supporting information for "Identification of CD44 as a key mediator of cell traction force generation in hyaluronic acid-rich extracellular matrices"

### SI MATERIALS AND METHODS

#### Temperature ramping for rheology measurements

The architecture of the collagen network depends sensitively on the rate of ramping temperature during polymerization step(1). To ensure that the mechanical properties of the ECM on rheometer were the same as the ECM used in our experiments, we made sure that the temperature ramping rate for collagen polymerization on the rheometer was the same as that in our *in vitro* experiments. To do that, we first measured the temperature ramp rate on the glass bottom dish during the routine polymerization steps in the incubator. An ultra-thin 10K thermistor (B3950 NTC, Adafruit) was taped onto the glass bottom and connected to a simple voltage divider circuit at 4℃. The system was then put in the incubator with 37℃ and the temperature was recorded until it plateaus at 37℃. Fig. S10 shows that it takes about 30 minutes for the system to reach steady state of 37℃ from 0℃. Since the rheometer platform only takes a linear temperature ramp function, we estimated that the temperature ramp rate through a linear fit to be 1.1℃/min (Fig. S10). We used the Peltier plate on the rheometer and the control within the rheometer to ensure that the ramping rate of collagen polymerization was 1.1℃/min.

#### Imaging device preparation

For all cell experiments, we used a 35mm glass bottom dish with No. 0 glass (D50-14-0-U, Matsunami) (Fig. S9). To ensure a consistent gel thickness across all the cell experiments, we created multiple (usually 4-9) wells using a 300μm-thick polydimethylsiloxane (PDMS) sheet. To prepare PDMS wells, we mixed 10 g silicone elastomer with 1 g curing agent from the SYLGARD 184 Silicone Elastomer Kit (Dow, USA) thoroughly. The mixture was degassed in vacuum for 20mins to remove air bubbles, followed by incubation at 60°C overnight for polymerization. Through-holes with a diameter of 2.5mm were created using a biopsy punch on the PDMS sheet. PDMS sheet and cover glass were bonded following plasma treatment with an oxygen plasma oven (PDC-001, Harrick Plasma) on “High” mode for 1min, followed by UV sterilization. To enhance attachment of collagen matrices to the glass/PDMS surfaces, surfaces were treated with 2$\mu$L of 1% polyethyleneimine (PEI) (P3143, Sigma-Aldrich) for 15 mins and then with 2$\mu$L of 0.1% glutaraldehyde (16019, EM Sciences) for 45 mins. The dishes were left in biohood overnight in sterile distilled water and were washed with PBS for three times before experiments.

#### Spheroid harvesting

To harvest spheroids, a 70-$\mu$m Falcon cell strainer (352350, Corning) was placed on top of a 50mL centrifuge tube (CLS430290, Corning), where spheroids were flushed out of the microwells using PBS. Spheroids were then gently collected from the cell strainer by washing with PBS. Spheroids in PBS were centrifuged at 1000g for 3mins. Supernatant was aspirated and spheroids were resuspended in 1mL complete medium.

#### Cell and spheroid embedding

Cell density was 50 cells/$\mu$L for 3D TFM experiments, and 400 cells/$\mu$L for single cell migration assays. 2.5$\mu$L of the cell embedded gel mixture was placed into each of the PDMS wells (2.5mm diameter and 300$\mu$m in depth) for polymerization in a 37℃ and 5% $CO_2$ incubator. To prevent cells from sinking to the bottom of the glass bottom dish during polymerization, the device was first placed upside-down for 5 min and then upright for 10mins. Subsequently, the dish was flipped twice at time points 15 min and 30 min respectively. Throughout the process, the glass bottom dish was contained within a second moist petri dish to prevent gel dryout. As such, it adopted a slow warming procedure for polymerization to obtain long and thick collagen fibers in contrast to the fast warming process described in our previous publication that yields thin and dense collagen fibers(1). After polymerization, 1.5mL complete medium was added to each glass bottom dish, and cells were incubated overnight at 37°C with 5% $CO_2$ overnight. 1 hour prior to the experiment, we replaced the culture medium and equilibrated cells with FluoroBrite™ DMEM (A1896701, Gibco) to minimize autofluorescence caused by phenol red. The medium was supplemented with 10% FBS (S11150, Atlanta Biologicals), 1X GlutaMAX™ (35050061, Gibco), 10mM HEPES (15630080, Gibco), 1% penicillin/streptomycin (15140122, Gibco), and 1mM sodium pyruvate (11360070, Gibco).

#### TFM imaging

1$\mu$m blue fluorescent carboxylated beads (F8814, ThermoFisher) were embedded and coupled to the matrix fibers as position markers at a bead density of ~530,000 beads/$\mu$L. Image stacks were acquired with a 20x magnification objective lens (NA = 0.45; LUCPlanFL N 20x, Olympus America) installed on an inverted epifluorescence microscope (IX81, Olympus America) and a 16-bit sCMOS camera (ORCA Flash 4.0 V3, Hamamatsu Photonics). Images of beads and cells measured

665.6 × 665.6 × 50 $\mu$m, with a voxel size of 0.325 × 0.325 × 1 $\mu$m ($x$, $y$, and $z$ respectively). Beads were imaged using a DAPI filter cube set (Chroma Technology, USA), and cells were imaged using the brightfield channel.

#### Traction force calculations

The general idea of traction force computation is to first calculate the matrix displacements using the measured ECM material properties, cell shape and presumed cell traction force, and then compare the computed matrix displacements with those of experimentally measured displacements. The program will adjust values of the cell traction force as well as ECM material properties iteratively until the theoretical displacements matches the experimental displacements. The resulted traction force is the measured traction force.

Computation of matrix displacements due to traction force was performed using a computation algorithm in COMSOL. The cell was modeled as an elliptical body with defined material properties, and the ECM was modeled as a cylinder much larger than the cell. The computation algorithm uses a previously developed fibrous nonlinear elastic model for ECMs and a predefined traction force to compute cell traction force. The initial input parameters to the computation algorithm were the material properties of the ECM measured from rheology experiments (this includes the initial differential shear modulus $K_o$, the onset for strain stiffening $\gamma_c$ and the strain stiffening slope $m$), cell traction force, and cell geometry. The theoretical matrix displacements are computed using the fibrous nonlinear elastic material model.

To match the theoretically calculated displacements with those of experiments, we implemented the Nelder-Mead method (a numerical computation package in MATLAB) to search for a minimum in a multi-parameter space. This method enhanced the search speed significantly from our previous least square optimization search method. Briefly, the adjustable parameters for the Nelder-Mead search are the material properties and the cell traction force. Logistically, our MATAB program was connected to COMSOL via the COMSOL LiveLink™ module. The MATLAB program does the minimum search, and the COMSOL algorithm does the displacement field computation, until the theoretical and experimental displacement field converges. The principal stress along the cell major axis is then used to calculate the traction force.

#### Neutralization of surface adhesion receptor molecules

To study the role of surface adhesion receptors CD44 (also known as HCAM) and $\beta$1-integrin in traction force generation, we neutralized these proteins using the following antibodies: 4$\mu$g/mL mouse monoclonal anti-HCAM (DF1485) (sc-7297L, Santa Cruz Biotechnology), and 4 $\mu$ g/mL mouse monoclonal anti-integrin $\beta$ 1 (P5D2) (sc-13590L, Santa Cruz Biotechnology). To validate that the effect of neutralization was not caused by the presence of an antibody itself, we performed the same experiment with 4$\mu$g/mL isotype control antibody (16-4714-82, Invitrogen).

#### Knockdown of CD44 and its validation

Knockdown of CD44 was achieved through lipotransfection with DsiRNA. Cells were plated on a 6-well plate at a density of $7 \times 10^5$cells/well in antibiotic-free complete media one day prior to transfection. On the next day, transfection mixtures containing RNAiMAX transfection reagent (13778075, Invitrogen), Opti-MEM (31985070, Gibco) and a pool of three DsiRNAs (IDT DNA Technologies) were prepared according to the manufacturer's protocol. A final DsiRNA concentration of 11nM was used in all experiments. A pool of three non-targeting DsiRNAs were used as a negative control. Cells were harvested 24hrs after transfection for experiments. For each experiment, we prepared extra wells of cells for immunostaining of CD44 (as described in the following section) following transfection to ensure successful knockdown before downstream experiments.

#### Immunofluorescence staining

To visualize CD44 and F-actin, we stained the cells with the according conjugated monoclonal antibodies. To do that, culture medium was aspirated and cells were washed three times in PBS at room temperature. Cells were fixed by a 15-min incubation in 0.1% glutaraldehyde/4% paraformaldehyde at room temperature. After fixation, cells were washed three times in PBS at room temperature, followed by permeabilization using 0.1% Triton-X100 for 30mins at room temperature. Then, cells were washed three times in PBS, and 1% bovine serum albumin (BSA) in PBS was used to block non-specific binding for 30mins at room temperature. Then, we applied 1:50 phalloidin-iFluor 488 reagent (ab176753, Abcam) and 1:50 anti-CD44 monoclonal antibody conjugated with AlexaFluor 647 (103018, BioLegend) on cells and incubated overnight at 4℃ inside a moist petri dish. To protect the fluorescently labeled cells from fading, cells were incubated with SlowFade™ Diamond Antifade mountant (S36963, Invitrogen) overnight at 4℃. For nuclear staining and protection from fading, SlowFade™ Diamond Antifade mountant with 4',6-diamidino-2-phenylindole (DAPI) (S36964, Invitrogen) was used instead.

#### Protein colocalization imaging and computation

To examine the colocalization between CD44 and F-actin, MDA-MB-231 cells were stained as described above. Cells were then imaged with a C-Apochromat 40x/1.20 W Corr M27 objective (Zeiss) on a LSM 710 Zeiss confocal microscope. Confocal *z* stacks of individual cells were acquired using a 488nm (for F-actin), and 633nm (for CD44) laser with a slice thickness of 1$\mu$m. To quantify colocalization, we computed the Spearman's rank correlation coefficient(2) slice by slice using an in-house MATLAB algorithm adopted from the ColocAnalyzer MATLAB package (Cambridge University) following background subtraction. Cells were segmented by the ImageJ plugin, U-Net, using a previously trained segmentation model(3).

#### Flow cytometry analysis of p-MLC

Collagen matrices containing cells were incubated with 1 mg/mL collagenase in the presence of 1x Halt™ phosphatase inhibitor cocktail (78420, Thermo Scientific) at 37℃ for 1hr. Cells were then centrifuged in a staining buffer with 2% FBS (26140079, Gibco) in PBS, followed by a 30-min incubation at room temperature in staining buffer containing 1x LIVE/DEAD fixable viability dye (L34990, Invitrogen) and 1x Halt™ phosphatase inhibitor cocktail (78420, Thermo Scientific). Cells were then washed and centrifuged, and fixed in 200uL 1.5% formaldehyde for 10mins at room temperature. After fixation, cells were centrifuged and washed in staining buffer, followed by permeabilization using 200uL ice-cold 99.8% methanol (322415, Sigma) for 15mins on ice. Cells were then centrifuged and washed twice in staining buffer. Cells were incubated with 2.5% normal goat serum (ab7481) for 10mins on ice to block non-specific binding. To stain for p-MLC, cells were incubated with 1:100 anti-p-MLC2 (Ser19) rabbit polyclonal antibody (3671, Cell Signaling Technology) for 15mins on ice, and 1:1000 goat anti-rabbit IgG H&L conjugated with Alexa Fluor 488 (ab150077) for 30mins at room temperature, with two washing steps in between. All samples were resuspended in staining buffer and kept on ice before analysis. All flow cytometry analyses were performed on the Attune NxT flow cytometer (ThermoFisher, USA). ~20,000 events were recorded for each sample. Unstained, single stained, and Y-27632 controls were used to determine the compensation matrix and negative gates for data analysis in FlowJo (Fig. S5B).

#### SUPPLEMENTARY FIGURES

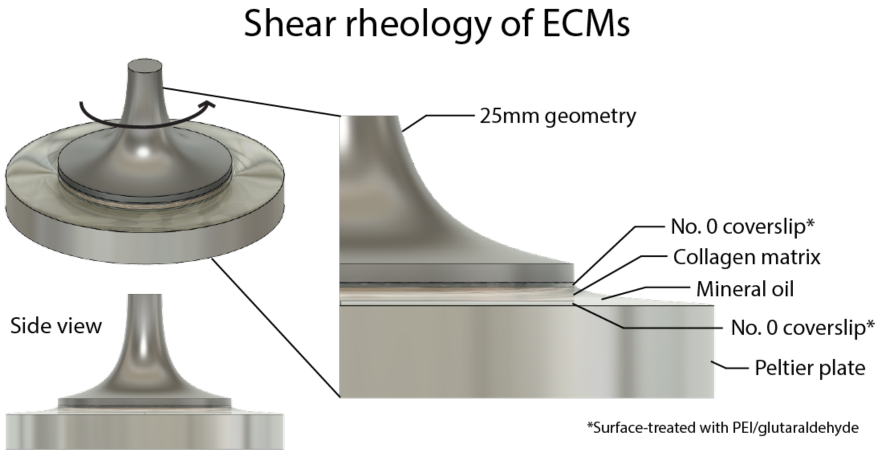


**Figure S1. 3D view of a parallel plate shear rheology setup.** A thin disc of 500$\mu$m-height and 25mm-diameter ECM was sandwiched between two coverslips that were glued onto the Peltier plate (bottom plate) and the 25mm geometry (top plate). Mineral oil was applied around the ECM to prevent evaporation during the rheology experiments. Collagen matrices were first polymerized on the Peltier plate with a controlled temperature ramp rate of 1.1℃/min to reproduce the polymerization rate of the ECM in our *in vitro* experiments. The 25mm circular geometry was used to apply oscillatory shear strain on the matrices. We note that two coverslips were used to sandwich the collagen to ensure proper anchoring of collagen to the surface. The coverslips were treated with PEI and glutaraldehyde prior to experiments.

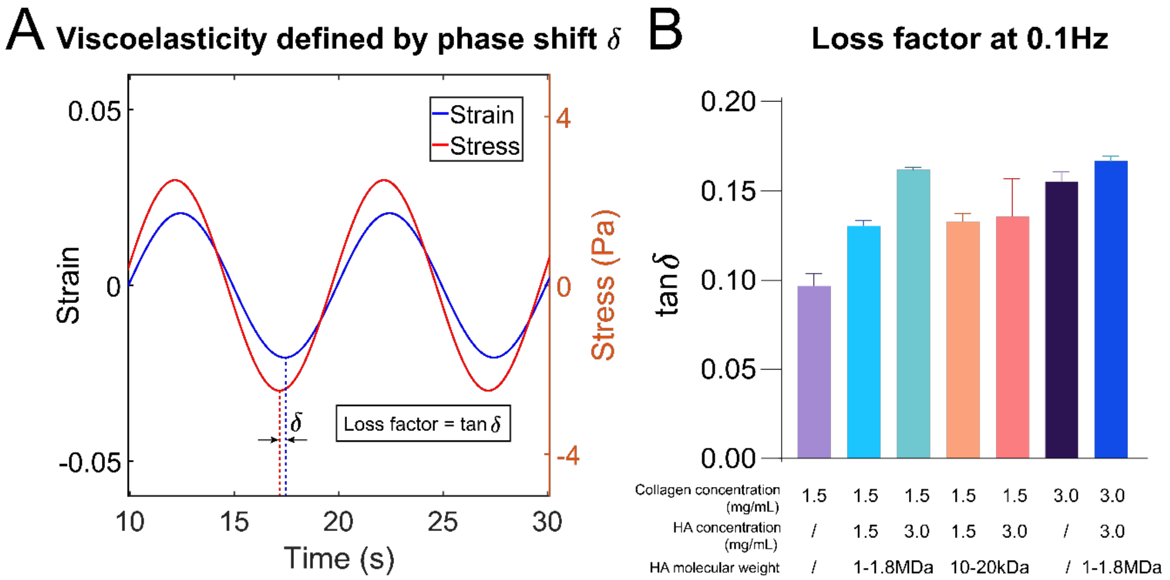


**Figure S2. Characterization of viscoelasticity of the ECMs using shear rheology. A.** To measure relative contribution of viscosity and elasticity of the ECMs, we measured the phase shift $\delta$ between the stress response to an oscillatory strain, and computed the loss factor ($\tan\delta$). A purely elastic ECM would have a loss factor of zero. The experimental data was fitted by sinusoidal functions. Note that loss factor increases with viscosity of the ECMs due to energy dissipation. **B.** Viscoelasticity of the ECM increases with collagen (Col) concentration, as well as HA (H and L) concentration. We have also tested whether the molecular weight of HA has an effect on viscoelasticity. Results show that cogels with high molecular weight HA (H) have higher viscoelasticity than cogels with low molecular weight HA (L). And viscoelasticity appears to be independent of concentration when LMW HA is introduced to the matrix. The numbers here represent concentration in mg/mL.

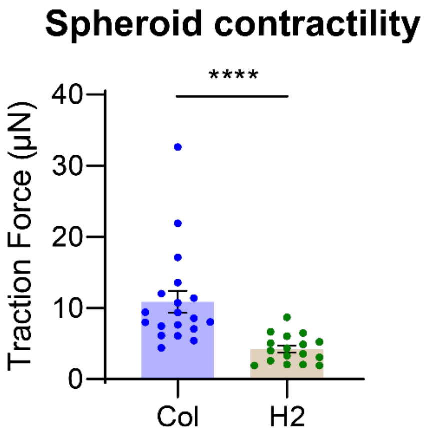


**Figure S3. Addition of HA significantly reduced tumor spheroid force generation.** TFM experiments using MDA-MB-231 spheroids show that traction force is significantly reduced in H2 cogel, compared to pure collagen (n = 19 for Col and n = 17 for H2). ****: p<0.0001, Student's t test.

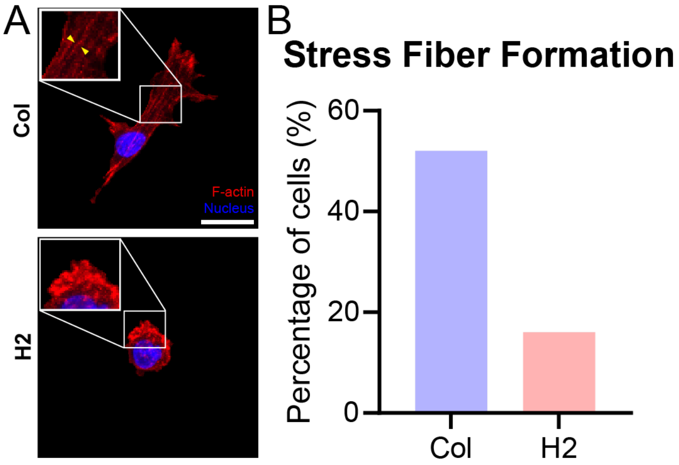


**Figure S4. Stress fiber formation in Col and H2. A.** We visually inspected cells throughout the confocal stacks and counted the percentage of cells that display filamentous actin structures (yellow arrows) (n=50 for both Col and H2). Scale bar = 20$\mu$m. **B.** Stress fibers are more prevalent in cells embedded in Col than in H2.

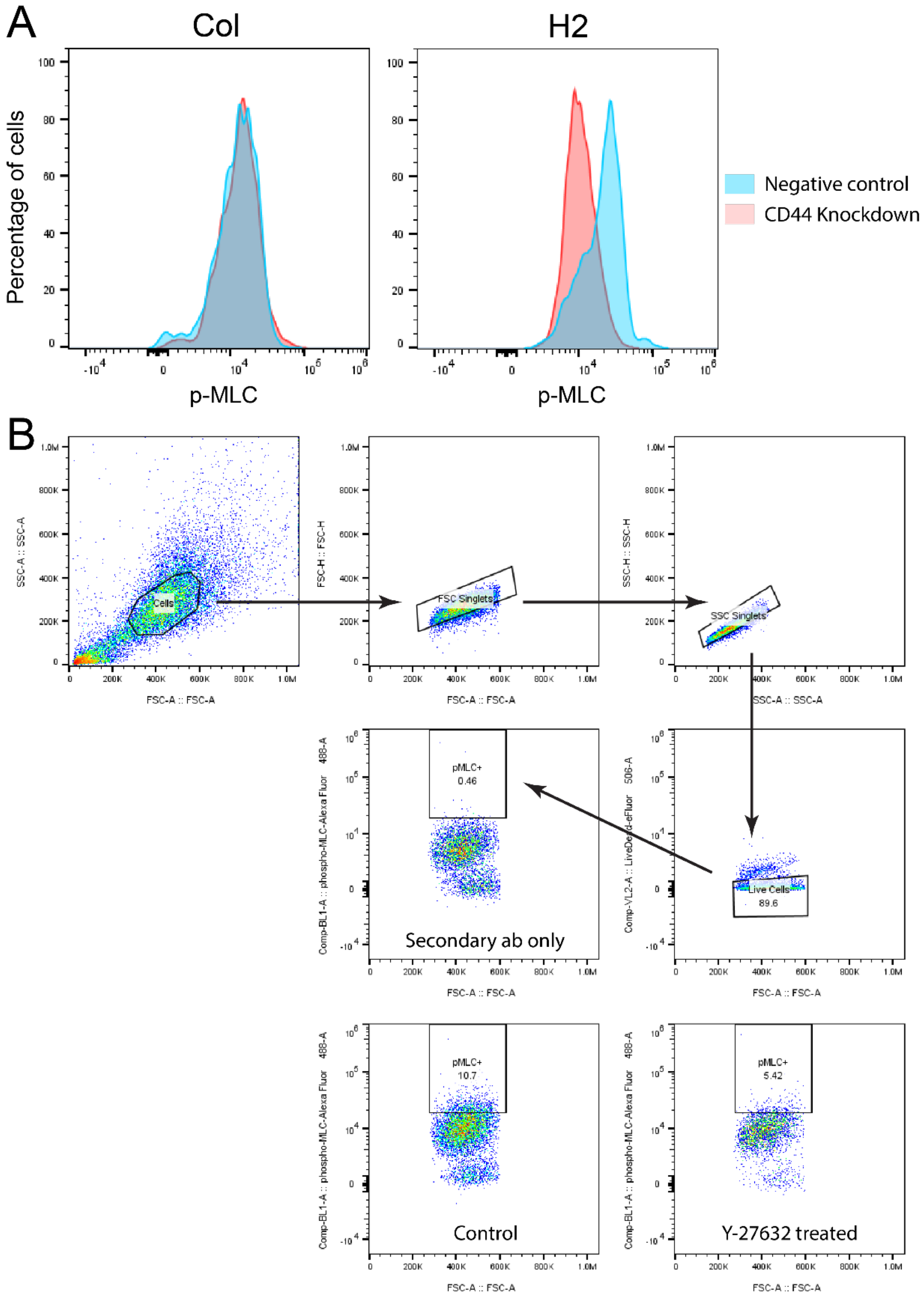


**Figure S5. The effect of CD44 knockdown on p-MLC expression. A.** Histograms of p-MLC expression of cells in Col and H2, with and without CD44 knockdown. p-MLC expression decreases in H2 when CD44 is knocked down, but not in Col, indicating that CD44 is required for actomyosin contractility in the presence of HA. **B.** Gating strategies for p-MLC. Y-27632-treated cells were used as negative controls for p-MLC gating.

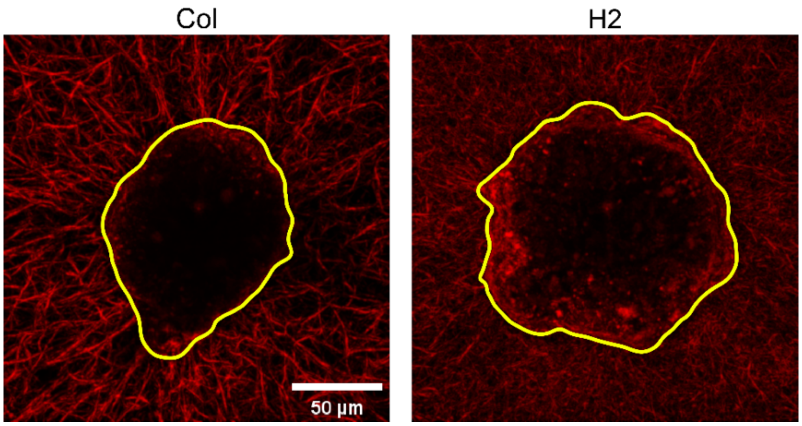


**Figure S6. Spheroids align the fiber network radially in Col, but not in H2.** 15$\mu$m-thick maximum intensity Z-projections of reflectance confocal images show that spheroids (outlined in yellow) contract and align the collagen fiber network (red) in pure collagen, but not in H2.

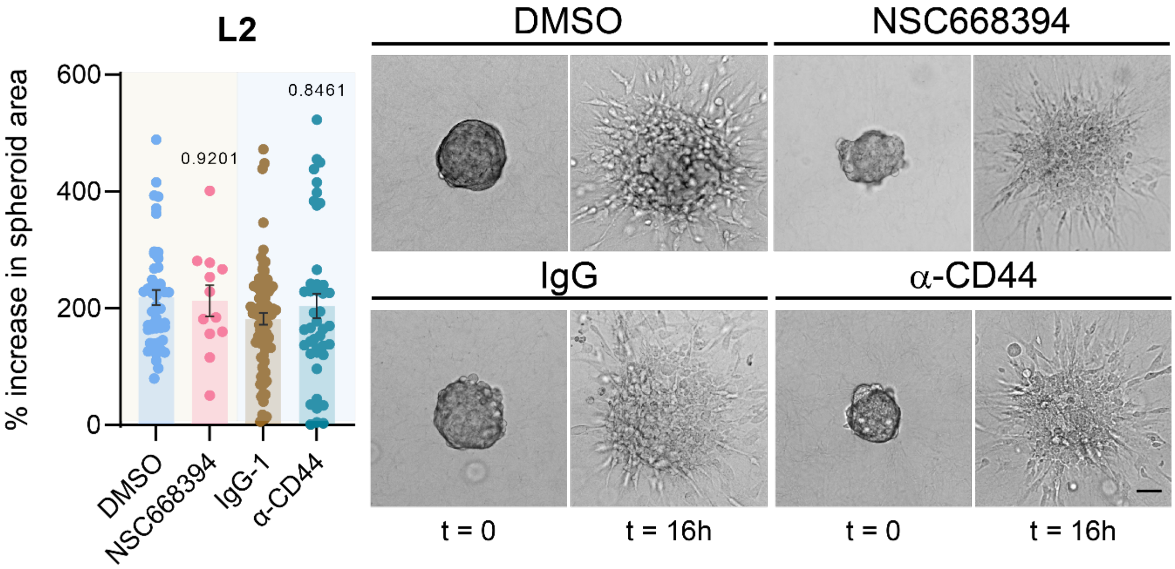


**Figure S7. Tumor invasion in LMW HA-rich environment is CD44-independent. A.** Spheroid invasion assay with ezrin inhibition (top row) and CD44 blockade. **B.** While tumor spheroids are more invasive in L2, compared to Col and H2, tumor invasion is insensitive to ezrin inhibition and CD44 blockade in LMW HA-rich environment. All statistical tests were Kruskal-Wallis tests. Scale bar = 50$\mu$m.

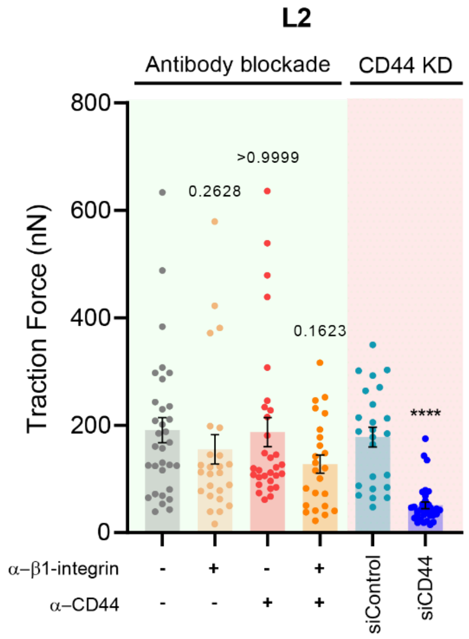


**Figure S8. Effect of CD44 modulation on traction force generation in LMW HA-rich environment.** CD44 knockdown was efficient in reducing traction force generation significantly in LMW HA-rich environment, while CD44 blockade did not render a significant reduction in cell traction force. All statistical tests were Kruskal-Wallis tests (****: $p<0.0001$).

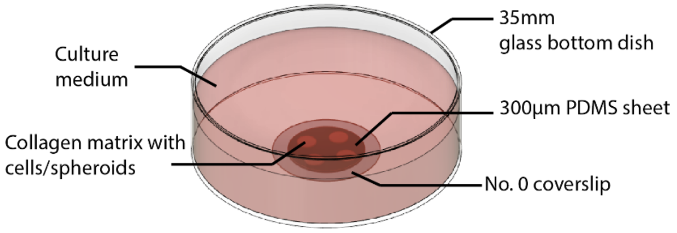


**Figure S9. Schematic of the imaging device.** A 300 $\mu$m-thick PDMS sheet was bonded onto the bottom of a No. 0 coverslip of the 35 mm glass bottom dish following an oxygen plasma treatment. The PDMS sheet was punched with multiple (usually 4-9) through-holes, which were used as microwells (2.5mm-radius and 300 $\mu$m-depth) to contain cell culture. The surface of the microwells was then treated with PEI and glutaraldehyde for crosslinking to collagen to avoid slippage during the experiment. Cell- or spheroid-embedded ECMs were then introduced into the microwells for imaging. The dish was filled with 1.5 mL culture medium during experiments. We note that the No. 0 cover slip was used to ensure a good imaging quality and minimal working distance since the thickness of it is about 100 $\mu$m.

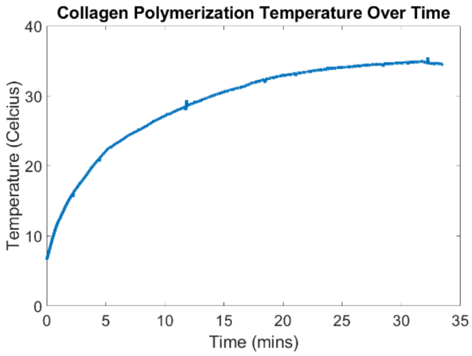


**Figure S10. Collagen polymerization temperature in the $CO_2$ incubator.** We monitored the polymerization temperature of the ECMs in the incubator and estimated the temperature ramp rate using a linear fit. Since the rheometer only takes a constant ramp rate, 1.1℃/min was used in all rheology experiments.

| Collagen concentration (mg/mL) | Molecular weight of HA | HA concentration | Initial shear modulus $K$ (Pa) | Critical strain for onset stiffening $\gamma_c$ | Strain-stiffening exponent $m$ |
|---|---|---|---|---|---|
| 1.5 mg/mL | / | / | 63.18±13.88 | 0.0474±0.0090 | 20.10±0.3755 |
| 3 mg/mL | / | / | 139.63±30.88 | 0.0172±0.0041 | 14.19±0.2427 |
| 1.5 mg/mL | High (1-1.8MDa) | 1.5 mg/mL | 42.66±1.36 | 0.0529±0.0049 | 18.01±0.1668 |
| 1.5 mg/mL | High (1-1.8MDa) | 3.0 mg/mL | 46.06±1.11 | 0.0750±0.0228 | 12.74±0.3868 |
| 1.5 mg/mL | Low (10-20KDa) | 1.5 mg/mL | 50.51±4.28 | 0.0194±0.0037 | 18.35±0.1898 |
| 1.5 mg/mL | Low (10-20KDa) | 3.0 mg/mL | 40.46±2.89 | 0.0281±0.0008 | 16.62±0.1865 |
| 3 mg/mL | High (1-1.8MDa) | 3.0 mg/mL | 76.37±22.43 | 0.02069±0.0051 | 12.99±0.7203 |

**Table S1**. Summary of mechanical properties of all the ECMs in this study, along with the computed initial shear modulus, critical strain for onset strain stiffening, and strain stiffening exponent.
